# Pesticide degradation capacity of a novel strain belonging to *Serratia sarumanii* (GBS19) with its genomic profile

**DOI:** 10.1101/2024.11.28.625876

**Authors:** Gülperi Alatassi, Ömür Baysal, Ragıp Soner Silme, Gülçin Pınar Örnek, Hakan Örnek, Ahmet Can

**Affiliations:** Molecular Microbiology Unit, Department of Molecular Biology and Genetics, Faculty of Science, Muğla Sıtkı Koçman University, Kötekli-Muğla, Turkey; Molecular Plant and Microbial Biosciences Research Unit (MPMB-RU), University of Worcester, Henwick Grove, Worcester, WR2 6AJ, United Kingdom; Center for Research and Practice in Biotechnology and Genetic Engineering, Istanbul University, Fatih-Istanbul, 34134, Turkey; Republic Turkey Ministry of Agriculture and Forestry, Directorate of Izmir Food Control Laboratory, Izmir, Turkey; Republic Turkey Ministry of Agriculture and Forestry, Directorate of Plant Protection Reseach Institute Bornova-Izmir, Turkey

**Keywords:** bioremediation, LC-MS, microbial degredation, *Serratia sarumanii* GBS19, Whole-genome sequence analysis

## Abstract

The use of pesticides in agricultural pest control and the potential for biological degradation of these chemicals are significant factors in environmental conservation. Sustainable pest control strategies should prioritize the preservation of non-target organisms and the maintenance of soil ecosystem dynamics. Initiation from this point, this study aimed to identify microorganisms capable of pesticide biodegradation commonly used in our region’s intensive agricultural fields. The microorganism with the broadest degradation spectrum was identified *in vitro* studies, and whole-genome sequence analyses indicated that some genomic regions were able to degrade pesticides, which was also confirmed with LC-MS analysis. Detailed genomic analyses linked to allele sequence definitions for ribosomal MLST have classified this strain as *Serratia sarumanii* under the *Serratia* genus. Moreover, our findings showed that the introduction of this microorganism can stimulate the degradation of certain pesticides in the environment while, interestingly, simultaneously enhancing the potency of low concentrations of some other pesticides through a synergistic effect. To validate of GBS19 biodegradation capacity, additional studies carried out on 25 pesticide active chemicals using LC-MS chromatographic analysis revealed that the *Serratia sarumanii* GBS19 strain is able to biodegrade some active compounds of various pesticides (fludioxonil, fenhexamid, pyrimethanil, and spirodiclofen) by up to 54%, 52%, 46%, and 23%, respectively. These results highlight the degrative properties of our prominent original strain as a gene pool which can be put into practice by transforming into competent strain using recombinant DNA technology.

## INTRODUCTION

The rapid growth of global industrialization and intensive agricultural practices driven by an expanding human population has led to significant pollution of ecosystems, including soil systems (Alori & Fawole, 2017). Pollutants present in soil exhibit immunotoxic, carcinogenic, and mutagenic properties, thereby altering the soil’s physical, chemical, and microbiological characteristics (Sales da Silva et al., 2020). Chemical pesticides are extensively used in various sectors of crop production to mitigate pest infestations, safeguard crop yields, and maintain product quality. A pesticide is any substance or mixture of substances designed to prevent, destroy, repel, or mitigate pests, including insects, mites, nematodes, weeds, and rodents. This category includes insecticides, herbicides, fungicides, and other agents used for pest control (Damalas & Eleftherohorinos, 2011; EPA, 2012). The application of pesticides has contributed positively by significantly enhancing crop and food productivity, and reducing the incidence of vector-borne diseases (Agrawal et al., 2010). Chemical pesticides can be classified in various ways, with the most common approach being based on their chemical composition. This classification method enables a systematic and scientific grouping of pesticides, facilitating the correlation between their structure, activity, toxicity, and degradation mechanisms, among other characteristics.

Worldwide, the annual pesticide consumption has stood at 3.69 million metric tons (FAO, 2024). Among these pesticides, insecticide use in the USA increased tenfold from 1945 to 2000 as reported by Sharma et al. (2019). However, a substantial proportion of these pesticides often fail to reach their intended targets due to degradation, volatilization, and leaching, resulting in significant ecological impacts (Chevillard et al., 2012). In typical agricultural practices, various groups of pesticides are frequently applied either simultaneously or consecutively, leading to interactions among them. A population residing in a contaminated site may experience selective pressure from the contamination, potentially resulting in higher resistance levels than a population of conspecifics inhabiting an uncontaminated site (Klerks et al., 2011).

Utilizing microorganisms’ capacity to eliminate pollutants in pesticide-contaminated environments has emerged as an alternative and viable approach. Bioremediation technology, which is an affordable, adaptable, and ecologically acceptable technique, comes to the fore in the strategy of using microorganisms to remove contaminants (Finley et al., 2010). Degradative enzymes that catalyze diverse reactions to transform pesticides into simpler molecules such as CO_2_, water, oxides, or mineral salts are found in microbial species, particularly bacteria (Doolotkeldieva et al., 2018) and fungi (Erguven, 2018). These degraded compounds can then be used as carbon, mineral, and energy sources (Senko et al., 2017). Many studies have focused on the metabolic and genetic underpinnings of microbial breakdown. Pesticide biodegradation-related genes were identified, described, and investigated. According to a previous study, various microorganisms have potential for biodegradation (Schroll et al., 2004).

The genus *Serratia* is classified within the family *Enterobacteriaceae* of the *Gammaproteobacteria*. *Serratia* spp. are ubiquitous microorganisms found in a wide range of environments, including water, soil, plants, and animals. Some *Serratia* spp. are classified as opportunistic human pathogens (Kurz et al., 2003). Atypical *Serratia* species (Stock et al., 2003), such as *S. ficaria* (Grimont et al., 1979), *S. fonticola* (Gavini et al., 1979), *S. odorifera* (Grimont et al., 1978), *S. plymuthica*, *S. rubidaea* (Ewing et al., 1973), *S. entomophila* (Grimont et al., 1988), and *S. quinivorans* (Ashelford et al., 2002), have been characterized as non-pathogenic (De Vleesschauwer, 2008). Unlike many other *Serratia* species, *S. plymuthica* has not concerned with pathogenicity in alternative animal models, such as *C. elegans* (Zachow et al., 2009).

*S. marcescens* is a well-established model organism for investigating biocontrol strategies. *In vitro* studies demonstrated that the antagonistic bacterium *S. marcescens* B2 suppressed the growth of *Botrytis* spp. and *Rhizoctonia solani* AG-1 IA (Someya et al., 2000; Someya et al., 2001; Someya et al., 2005). *S. marcescens* strain B2 reduced the severity of rice blast disease, resulting in smaller lesion sizes caused by the pathogen (Someya et al., 2002). Additionally, previous studies identified *Serratia marcescens* strains (*MEW06; NCIM 2919;* DT-1P) that are able to degregade pesticide and petroleum derivatives as carbon sources (Wang et al., 2018; Grewal et al., 2016; Bidlan and Manonamani, 2002).

The aim of this study is to determine the presence of microorganisms that degrade pesticides in soil samples collected from production areas. The samples were collected from production fields to reveal their bioremediation potential. In addition, it was investigated in which pesticide groups this microorganism could be used more effectively. For this purpose, whole-genome sequence analyses of the detected bacteria were performed, and the genes responsible for pesticide degradation were determined. Moreover, advanced chromatographic analyses were performed to investigate the potential for the degradation of chemical compounds belonging to different pesticide groups by this microorganism.

## EXPERIMENTAL PROCEDURES

### Collecting soil samples for bacterial isolation

Soil samples from various greenhouses to be analyzed for pesticide bioremediation were collected from Ortaca district of Muğla province. A randomly selected 5 g of soil sample was diluted with 15 ml of distilled water and 5 µl of cymoxanil fungicide to inhibit fungal growth and contamination. The solutions were shaken at 90 rpm for three hours before filtering using Whatmann filter paper. One cubic centimeter of the filtered solutions was separately added to the NB liquid medium and three days incubated at 27°C. The visual colonies that appeared on petri dishes were considered to be able to degrade various pesticides (the pesticide list is given below). The colonies were then kept in pure culture at +4°C.

Colony formation and visual appearance on culture media were typical properties of the *Serratia* genus; therefore, we carried out some typical biochemical tests to understand the genus of bacteria in our laboratory conditions.

### In vitro biodegradation analysis

The concentrations of the whole pesticides tested in culture media for performing the bacterial colonies are shown in Supplementary table S1 and Supplementary table S2.

#### Insecticide

Abamectin (18 g/l) which is the active compound of commercial pesticide, was used for insecticide biodegradation. Mediums were prepared according to the recommended concentrations of the insecticides (2,5 ml/10 l, 5 ml/10 l, 7,5 ml/10 l). Adjusted insecticide concentrations were mixed with homogenized sterilized aqueous agar. Inoculated Petri dishes were cultured at 27°C for 5 days. The appeared colonies were transferred to Petri dishes containing higher insecticide concentrations.

#### Fungicide

Penconazole (100 g/l) is another active compound tested for biodegradation. The following concentrations (2,5 ml/10 l, 3,5 ml/10 l, and 5 ml/10 l) were used considering recommended doses. Adjusted fungicide concentrations were mixed with homogenized sterilized aqueous agar. Inoculated Petri dishes were cultured at 27°C for 5 days. The appeared colonies were transferred to Petri dishes containing higher fungicide concentrations, which were administration levels.

#### Herbicide

Glyphosate (480 g/L) is also other active compound. Adjusted Herbicide concentrations (30 ml/10 l, 60 ml/10 l, 100 ml/10 l). The appeared colonies were selected as described above.

### Selection of a pesticide-biodegrading bacterial strain

Bacterial strains exposed to various concentrations of different types of pesticides that enabled them to survive due to their carbon preference indicated their biodegradation capacity when the pesticide concentrations increased in culture media. The bacterial colonies surviving in the highest concentration of whole tested pesticides were selected for cross-tests (the colonies that are able to survive at the highest concentration of any pesticide were transferred from one to another petri dishes containg different pesticide groups). Finally, the cross-tests indicated that the bacterial strain degraded the entire types of tested pesticides (Supplementary table S1 and table S2). Our study involved genomic characterization and LC-MS analysis on this selected bacterial strain.

### Whole-genome sequencing of bacterial strain

The bacterial samples were cultured in NB medium for 1 day, followed by centrifugation. DNA was purified from the pellet sections using the CTAB method (Nishiguchi et al., 2002). The purity of the extracted genomic DNA was assessed using 1% agarose gel. The purified DNA was used to construct a library with blunt triple junctions consisting of fragments of approximately 500 base pairs in length. Subsequently, the library was subjected to sequencing using an Illumina Hi-Seq 2500 sequencing system.

We performed adapter trimming and quality filtering on the reads using Trim Galore version 0.6.7 (Krueger, 2020), which incorporates Cutadapt version 3.5 (Martin, 2011). The - q parameter was set to 30, and we used the --paired option. The resulting cleaned read pairs served as input for de novo assembly using SPAdes version 3.15.5 (Bankevich et al., 2012). The resulting scaffolds and contig were re-ordered against the reference genome of strain AR_0027 (NCBI acc. num; CP028702.1). The final assembly was evaluated using QUAST 4.5. Annotation was performed using the PGAP pipeline and RAST (Rapid annotation using subsystem technology) (Brettin et al., 2015). The genomic sequence was also annotated using BLAST and compared with the UNiprot protein database (Tatusova et al., 2016).

The total number of forward and reverse primer sequences were 10,167,833 bp. The final genome assembly comprised 658 contig with a total length of 10,167,833 bp. The longest segment spanned 282,998 bp, and the N50 value was 35,605 (Supplementary data S1). To identify the predicted genes, the assembled genome was subjected to Quast v. 5.0.2 analysis (Gurevich et al., 2013). Subsequently, PATRIC (Wattam et al., 2017) was used for further analysis.

Protein sequences were annotated with Enzyme Commission (EC) numbers (Schomburg et al., 2004), Gene Ontology (GO) terms (Ashburner et al., 2000), and KEGG pathways (Kanehisa et al., 2016) using genome annotation pipelines (e.g., RAST and PROKKA) (Aziz et al., 2008; Seemann, 2014). The antibiotic production and bacteriocin genes were identified using the SEED viewer server (Aziz et al., 2012). An in-depth analysis of the antibacterial compounds produced by GBS19 was conducted using AntiSMASH (version 6.0) to predict gene clusters involved in the production of secondary metabolites (Blin et al., 2021).

From the resulting contig, the 16S rRNA sequence was extracted and compared with reference genomes available in the databases. A phylogenetic tree was constructed using the neighbor-joining method with related sequences obtained from the BLAST output of NCBI (2007).

Gene profiles of the bacterial samples were created by examining the sequence data in the KEGG database, and species determination was performed by extracting 16S sequences from the sequence data. The entire genome of the bacterium (GSB19 NCBI acc. num; PRJNA784190 https://www.ncbi.nlm.nih.gov/nuccore/JAKDDT000000000.1) was mapped against the reference genome (Reference genome: AR_0027 NCBI acc. num; CP028702.1).

From the resulting contig, 16S rRNA sequence was extracted and compared with reference genomes available in the databases. A phylogenetic tree was constructed using the neighbor-joining method with related sequences obtained from the BLAST output of NCBI. Additionally, the GBS19 gbk file was analysed for the average nucleotide identity (ANI) and DNA-DNA hybridisation (dDDH) using The Type (Strain) Genome Server (TYGS) (Meier-Kolthoff & Göker, 2019). Additionally, the fasta sequence of the whole-genome was submitted to PubMLST (https://pubmlst.org/), and allelic variation analyses were performed. Furthermore, the pathogenic gene regions were investigated using Pathogen IslandViewer 4.0 (Bertelli et al., 2017).

### Degradation pathways of *S. sarumanii* GBS19

The strain of *Serratia sarumanii* GBS19 was identified as the bacteria that metabolized pesticides using the full genome sequence. The strain’s position in the family tree, the enzymes it encodes, and the degradation processes linked to whole-genome sequencing analysis. Degradation routes and xenobiotic degradation pathways were assessed among the strain’s related pathways and encoded enzymes.

### *S. sarumanii* GBS19 genes and critical nodes in degradation pathways

Based on the enzymes expressed by the strain, enzymes that catalyze metabolic processes at crucial points of the breakdown pathways with which the *S. sarumanii* GBS19 strain is involved were evaluated. Using the genes of *Serratia marcescens* AR_0027 (acc. num; CP028702.1) as a guide, the NCBI blast software was used to determine whether the *S. sarumanii* GBS19 strain encodes these enzymes.

### Metabolomics prediction of GBS19

The fasta aminoacid sequences of the whole-genome data were submitted to PIFAR-Pred and PGPT-Pred of PLaBAse (https://plabase.cs.uni-tuebingen.de/pb/plabase.php) using Python modules (Patz et al., 2021; Patz et al., 2024; Martínez-García et al., 2016; Ashrafi et al., 2022), and the results were depicted with KronaTools (Ondov et al., 2011).

### LC-MS analysis for the degradation of active substances

Twenty-five pesticide active components were tested using *S. sarumanii* GBS 19 strain for biodegradation analysis using LC-MS/MS (TSQ Quantiva/ Ultimate 3000 (Pump)/ Ultimate 3000 (Autosempler Optiplex 9020/Sogevac). Administration dosages prepared with these active components were introduced with the *S. sarumanii* GBS19 strain for 72 h, and the biodegradation of these active compounds was measured. Calibration curves were constructed for all active substances, which showed a bioremediation effect on pesticides (Supplementary data S2, Supplementary table S3, Supplementary table S4).

A sensitive scale was used to weigh the powder formulations, which were then placed in 5 l sterile bottles containing up to half water and stirred with a magnetic stirrer for five min. All preparations containing active substances were prepared in the specified quantities. Depending on the application dose, various dilution factors have been established for each active component. Supplementary table S4 provides information on the active components of the pesticides used in LC-MS/MS analyses and provides details on the agricultural pests and diseases targeted by the active ingredients. Analysis was also performed with LC-MS and GC-MS depending on the active ingredient of the pesticide and the degradation amounts were determined according to the best response value. Detailed information on GC-MS analysis was given in Supplementary table S4.

## RESULTS

Bacterial groups in labeled tubes were examined for microbial biodegradation after shaking incubation. The isolates were analyzed for the biodegradation of commercial pesticides such as insecticides, herbicides, and fungicides.

The preliminary phenotypic characterization tests indicated that the bacterial isolate belonged to the *Serratia* genus (Suplementary table S5, Supplementary figure S1). To identify the bacteria, we continued our studies with genome-scale analysis.

*In vitro* tests showed that GBS19 can degrade various pesticides because the colonies were viable and visible on culture media even at high concentrations. Afterwards, we carried out LC-MS analysis to observe the degradation capacity considering active compound quantity.

### Whole-genome sequence analysis and identification

The whole-genome sequence and its encoding enzymes with phylogenetic tree analyses have been extensively investigated (Supplementary data S3). We submitted it to NCBI with the accession number PRJNA784190. Genetical identification was illustrated using PROKSEE and whole-genome data, and the distribution of CDS, tRNA genes, rRNA genes, ORF regions, GC skew +/−, CARD genes, and Phage integration regions in the GBS19 genome is shown in Figure 1a (Supplementary data S4). The Average Nucleotide Identity (ANI) was depicted considering reference genome *S. sarumanii* K-M0706. Even 16S rDNA analysis indicated that our strain GBS19 was genetically close to *S. nematodiphila* DSM 21420 (Figure 1c), DNA-DNA hybridisation (dDDH) proved that our strain belongs to *S. sarumanii* genus (Supplementary data S5) (Figure 1b), which was confirmed by ANI analysis (Figure 1d). To validate our genomic identification results, we confirmed our data using PubMLST database. Additionally, detailed genomic analyses linked to allele sequence definitions for ribosomal MLST have shown that this strain belongs to *Serratia sarumanii* under the *Serratia* genus, according to Jolley et al. (2012) using rMLST (Supplementary data S6) (Figure 2). Furthermore, 7 housekeeping genes with allelic variations were determined using MLST analysis by PubMLST after submitting genomic data to the server (Jolley et al., 2012) (Supplementary data S7).

**Figure 1.**
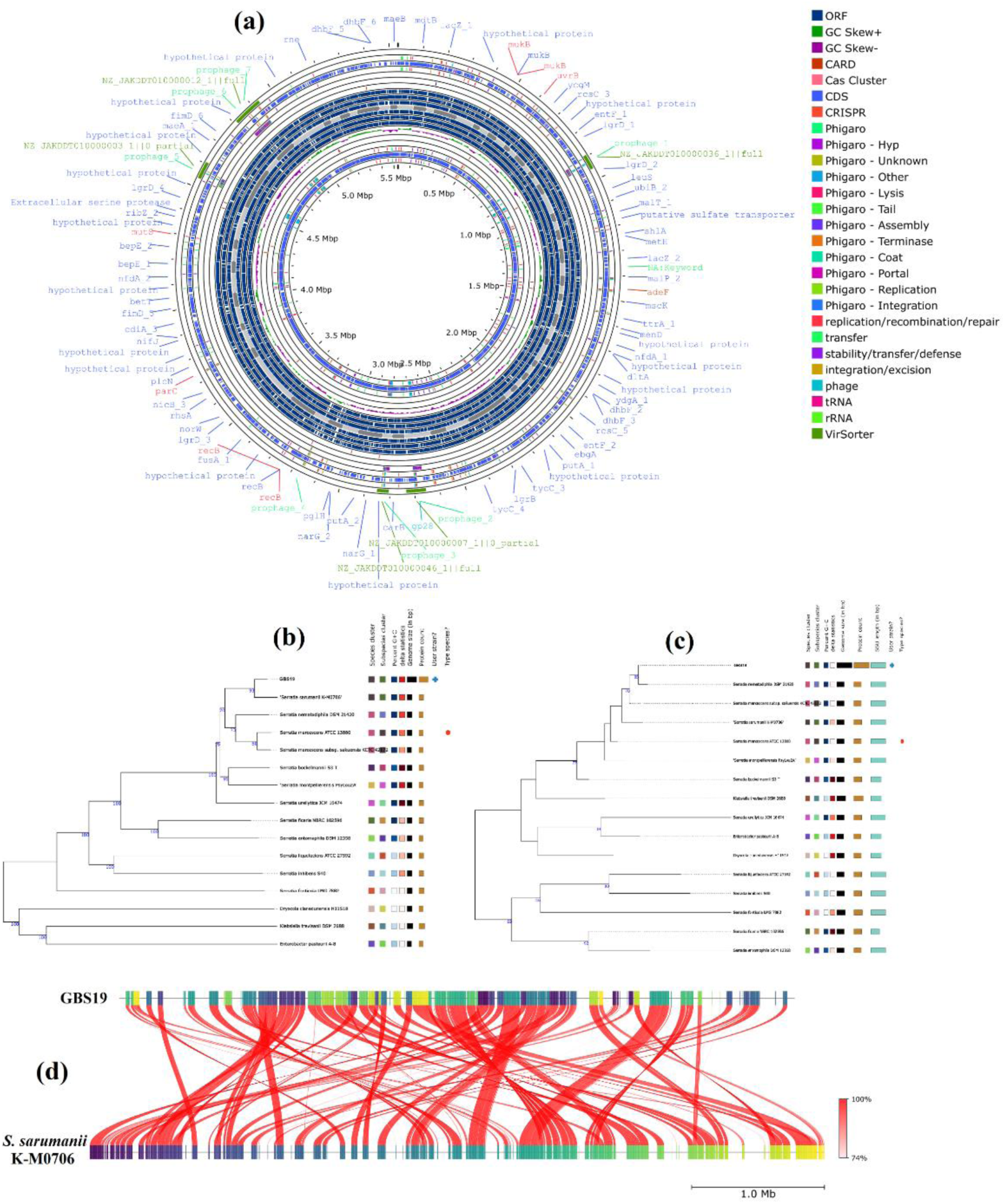
The figure shows genetical identification depending on whole-genome sequence data of GBS19. **(a)** The illustration shows a circular graphical display of the distribution of the genome annotations: CDS, tRNA genes, rRNA genes, ORF regions, GC skew +/−, CARD genes, and Phage integration regions in GBS19 genome. (**b**) The data depicts DNA-DNA hybridisation (dDDH) using the TYGS database. “**+**” denotes our GBS19 strain, which is genetically similar to *S. sarumanii* K-M0706. (**c**) The figure shows the 16S rDNA results obtained using the TYGS database. “**+**” denotes our GBS19 strain which is genetically similar to *S. nematodiphila* DSM 21420. (**d**) Whole-genome average nucleotide identity (ANI) comparison between reference *S. sarumanii* K-M0706 and GBS19.

**Figure 2.**
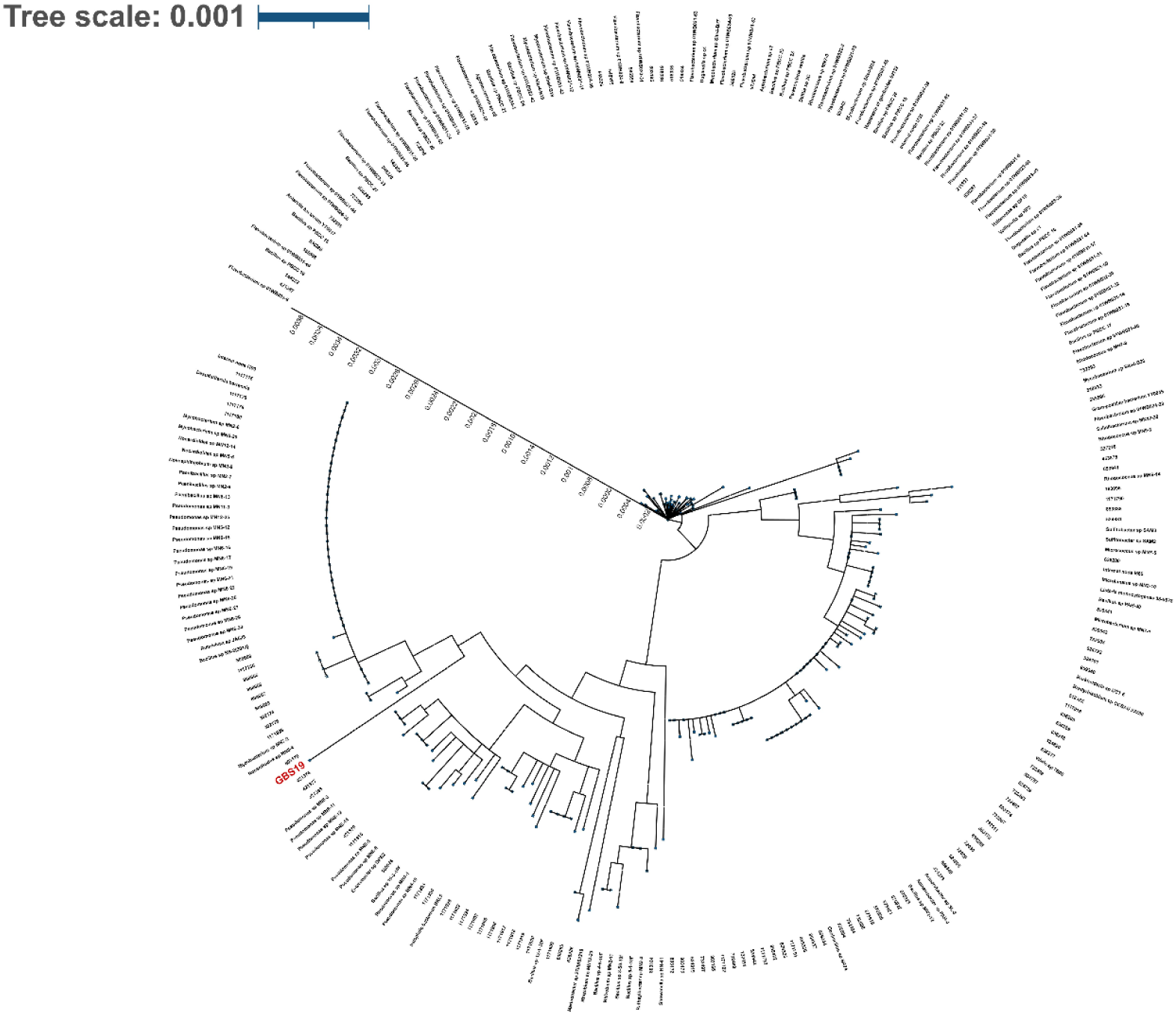
Ribosomal MLST analyses output obtained from PubMLST database and the phylogenetic tree was constructed considering allelic variations of GBS19 (shown in red) using the ITOL tool.

The number of *S. sarumanii* GBS19 genes related to molecular functions and cellular components (Figure 3a), biological processes (Figure 3b), and comprehensive antibiotic resistance (Figure 3c) are presented. In accordance with the subsystem feature count of the genes in the genome linked to the metabolism of aromatic compounds, 71 genes were identified in the GBS19 strain (Supplementary figure S2). Supplementary table S6, S7 and S8 show the encoding genes linked to degradation pathways in *S. sarumanii* GBS19 genome.

**Fig 3.**
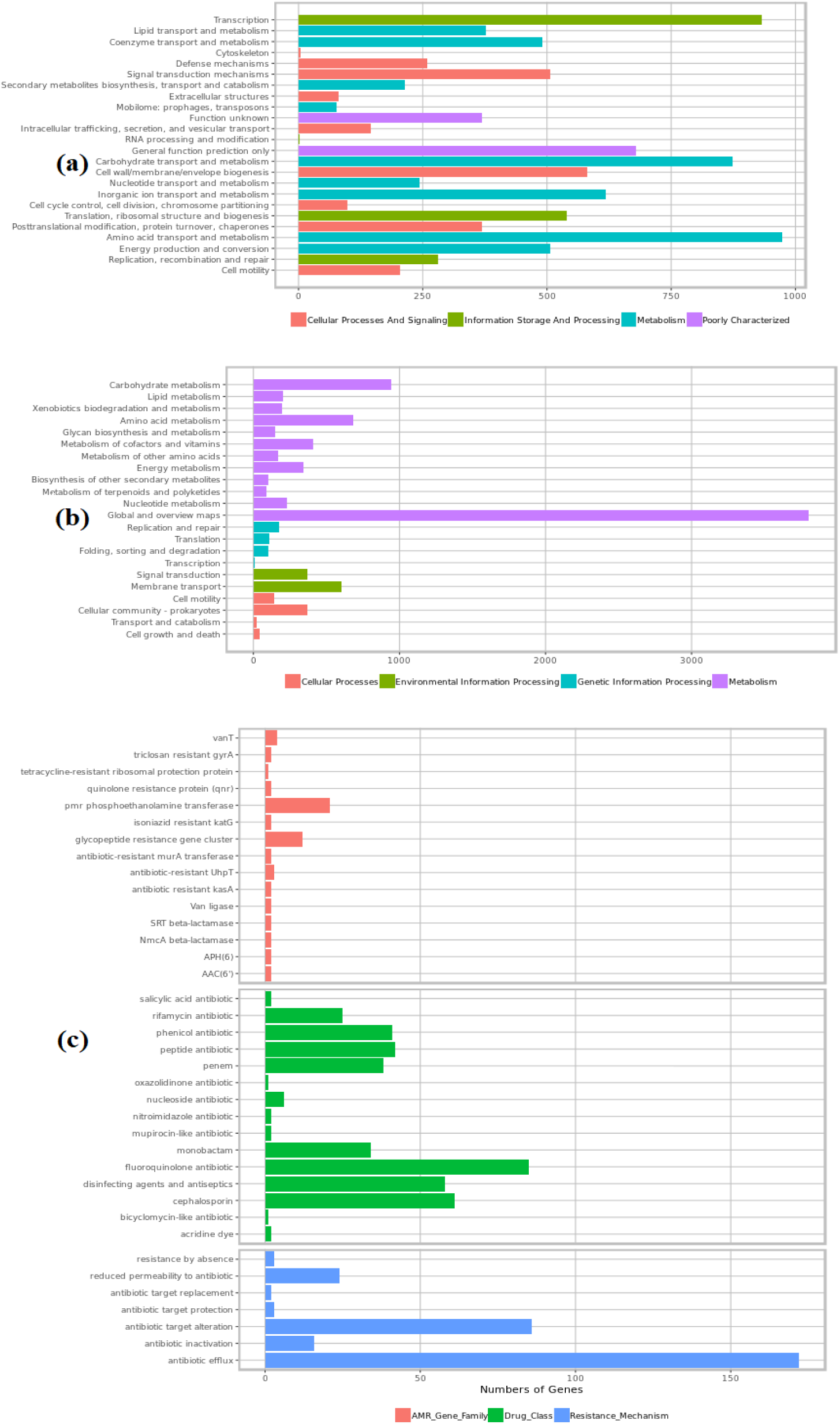
Number of the *S. sarumanii* GBS19 genes related to **(a)** molecular functions and cellular components, **(b)** biological processes, **(c)** comprehensive antibiotic resistance database outputs according to genomic data submitted from SEEDviewer 2.0.

*S. sarumanii* GBS19 strain’s molecular function, cellular component, and biological process percentages were determined using the entire genome sequence information (Figure 3). We searched for links between the enzyme information encoded by *S. sarumanii* GBS19 strain and mechanisms related to enzymatic pathways. The KEGG database mapping metabolic pathways and enzyme classification were determined according to Kanehisa et al. (2016) (Supplementary data S8).

The whole-genome sequence of GBS19 was submitted to the antiSMASH database and compared with the other available gram-negative and gram-positive prokaryotic genomes. The results showed a similarity between the antimicrobial metabolites, as depicted in Figure 4 (Supplementary data S9). Furthermore, Pathogen IslandViewer 4.0 results revealed that there were no pathogenic regions in our strain GBS19 (Supplementary figure S3).

**Fig. 4.**
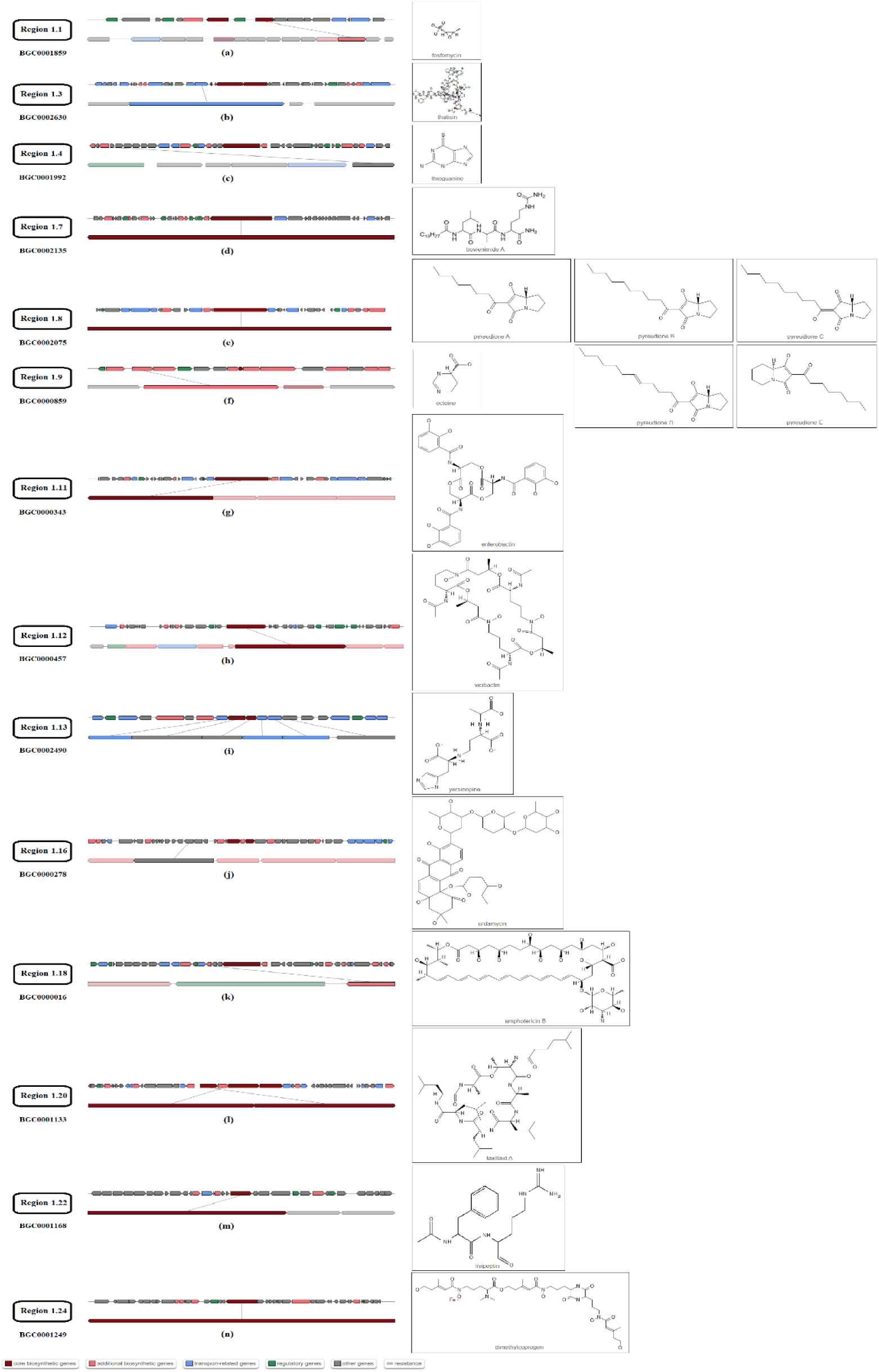
Antimicrobial metabolites produced by GBS19 identified using the antiSMASH database according to whole-genome sequence data. Genes encoding proteins responsible for biosynthesis of (**a**) fosfomycin, (**b**) thatisin, (**c**) thioguanine, (**d**) bovienimide A, (**e**) pyreudione A, B, C, D, E, (**f**) ectoine, (**g**) enterobactin, (**h**) vicibactin, (**i**) yersinopine, (**j**) urdamycin, (**k**) amphotericin B, (**l**) taxlllaid A, (**m**) livipeptin, and (**n**) dimethylcoprogen. Repetitive metabolites and metabolites below 0.20 MIBiG comparison value are ignored and not considered.

As depicted in supplementary data 1, the genes encoding enzymes were identified and listed. All genes related to the degradation of aromatic compounds showed that the GBS19 can degrade hydrocarbons and various chemicals.

### Predictive metabolomics profiling

Predictive metabolomics profiling of GBS19 using the PLaBAse database indicated a variety of disparities in the produced metabolites. These genes mainly encode toxins, exopolysaccharides (EPS), adhesion and e.g. related compounds (Figure 5a and 5b).

**Fig. 5.**
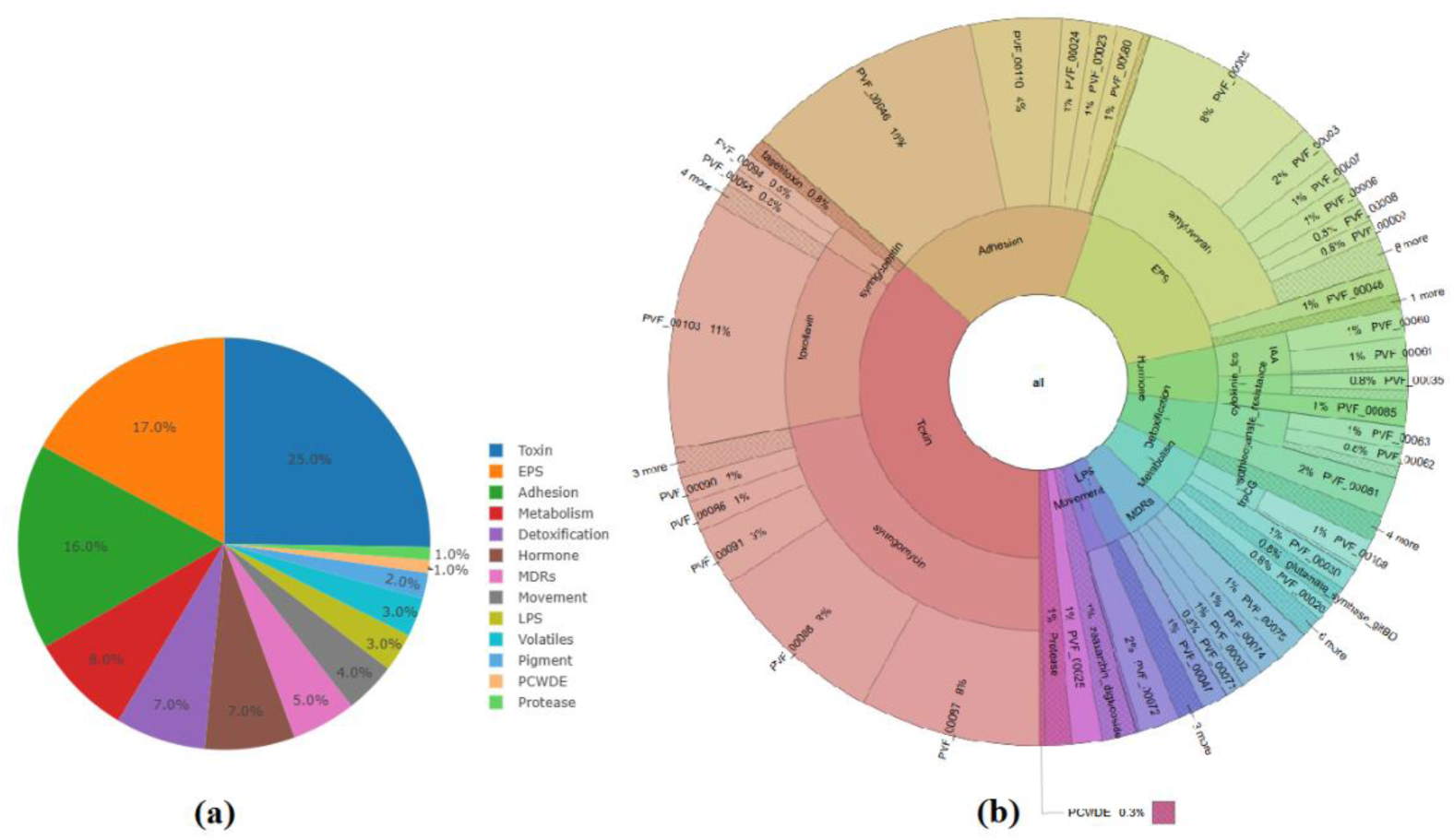
**a)** Pie chart and **b)** Krona chart of predicted metabolites produced by GBS19.

### LC-MS analysis results for the pesticide-active substance degradation

Among all tested pesticide groups shown in the Supplementary table S4, GBS19 was found to significantly degrade one pesticide from Group 2, three from Group 3, one from Group 5, three from Group 6, one from Group 7, and two from Group 8. The highest degradation rates were observed for Pyrimethanil, Fenhexamid, Spirodiclofen, and Fludioxonil (Supplementary table S4).

**Table 1.** LC-MS analysis of the active ingredients. Values of +/− denote the standard deviation of 3 replicates. Asterisks indicate the significance of the measurements.

| Groups | Pesticide Active Ingredients | Standard | GBS19 (+) 72 h | GBS19 (-) 72 h |
| --- | --- | --- | --- | --- |
| 2. Group | Pyrimethanil | 957±0.02 | **435\*\*±0.01** | 899±0.02 |
| 3. Group | Fenhexamid | 2829±0.03 | **1458\*\*±0.02** | 1771±0.01 |
|  | Indoxacarb | 131±0.03 | 111*±0.02 | 217±0.01 |
|  | Spirodiclofen | 420±0.02 | **98\*\*\*±0.03** | 306±0.03 |
| 5. Group | Metrafenon | 426±0.01 | 302*±0.03 | 354±0.01 |
| 6. Group | Azoxystrobin | 613±0.01 | 379*±0.03 | 416±0.01 |
|  | Difenoconazole | 324±0.03 | 319*±0.02 | 411±0.01 |
|  | Fludioxonil | 477±0.03 | **255\*\*±0.03** | 334±0.02 |
| 7. Group | Gamma Cyhalothrin | 42±0.02 | 40*±0.03 | 48±0.01 |
| 8. Group | Boscalid | 5503±0.01 | 4199*±0.01 | 4388±0.03 |
|  | Spirotetramat | 322±0.02 | 199*±0.01 | 287±0.03 |

## DISCUSSION

Worldwide, annual pesticide consumption shows an ascending trend, depending on agricultural production and consumption. Nevertheless, the environmental behavior and ecotoxicological impacts of many pesticides remain poorly understood (Carvalho, 2017). Post-application, a considerable number of pesticides penetrate the soil, where processes such as mineral adsorption and microbial degradation act as natural filters, protecting groundwater and surface water from contamination (Sun et al., 2010; Keesstra et al., 2012). The ability of microorganisms to remove pollutants by disintegration through various enzymatic reactions in areas contaminated with pesticides is an alternative and promising method. The methods of microorganisms to remove pollutants are provided via a technology known as bioremediation, which has emerged as an economical and environmentally friendly approach (Finley et al., 2010). The underlying principle of bioremediation is to decompose toxic contaminants through biodegradation and transform them into less toxic or non-toxic elements or compounds (Strong & Burges, 2008). Silva et al. (2019) reported that only 17% of 317 agricultural topsoil samples from the European Union were free of pesticide residues. While 25% of the soils were contaminated with individual pesticide residues, 58% contained pesticide mixtures at medium and high concentrations ranging from 0.02 to 0.04 mg kg^−1^ and 0.31 to 0.41 mg kg^−1^, respectively.

Despite their biodegradability under ideal laboratory conditions, numerous organic pollutants persist in soil environments (e.g., 2,4-dichlorophenoxyacetic acid) (Nowak et al., 2011; Fenner et al., 2013). Numerous studies have highlighted the adverse environmental impacts of conventional pesticides. In addition to harming non-target organisms within the ecosystem, these chemical pesticides can also migrate beyond the application site (Kalia & Gosal, 2011). Since the side effects of pesticides on human and animal health have shown by different studies (Bassil et al., 2007; Hayes et al., 2010), there is a great interest among researchers and farmers for more environmentally friendly approaches (Anonymous, 2024).

The population dynamism of microflora may differ at genomic level according to abiotic and biotic stress. Bacteria become resistant to various stresses, including antibiotics and pesticides, because of their evolved genes. This approach changes the genome stability and the survival of bacteria (Davies & Davies, 2010). Pesticide-degrading enzymes are encoded by the genes of bacteria present on chromosomes, transposons, and plasmids (Bahig et al., 2008).

The bacterium that we found to metabolize pesticides belongs to *Serratia sarumanii*, according to the genomic analyses, ribosomal MLST and MLST (considering 7 housekeeping genes). The identification analysis of the strain revealed that it was closely matched by 81% with *Serratia sarumanii* and 10% with *Serratia nematodiphila* (Supplementary data S6 and Supplementary data S7). In our research, we submitted this strain into the database as GBS19 and registered its whole-genome sequence as a novel strain in the NCBI (Acc. Num: PRJNA784190). Furthermore, in the identification of bacteria, there may be situations in which using 16sRNA analysis alone, in addition to phenotypic characterization, may give misleading results, and even MALDI-TOF MS analyses could be insufficient to respond to new taxonomic distinctions at the subspecies level, and we have also experienced this situation during our studies.

In fact, when we carried out genomic studies after some preliminary information at the genomic level in phenotypic characterization, it was determined that the isolate we had was *Serratia nematodiphila* in 16sRNA analysis, but in genome-to-genome comparison, it was identified as *S. sarumanii,* and the result causing an indecision was encountered. In this case, we expanded our analysis using rMLST and MLST database. Only then were we able to identify that the isolate we had belonged to *S. sarumanii,* as found by genome-genome hybridization analysis. Our data show that when identifying a bacterium at the genomic level, examining genetic allelic variations and characterizing them via housekeeping genes yield more reliable results. We found it appropriate to highlight this output as it will help in further studies on the identification of various bacterial strains.

As sequence analysis information in databases accumulates over time, it is possible that populations belonging to a certain group within the strains currently defined as *S. marcencens* may move out of the present group and become involved into another subspecies that is the closest to the group that can be classified as *Serratia sarumanii* (Klages et al., 2024). In fact, since our isolate was isolated from a soil where agricultural production was carried out, it showed a close relationship to *S. nematodiphila*, the closest to it, based on genetic results, rather than being a human pathogen. Our strain was isolated from soil; therefore, encountering the strain in human and its potential to become a human pathogen seems to be low probability. We also confirmed the nonpathogenic properties of our strain using Pathogen IslandViewer 4.0 (Bertelli et al., 2017), although it was first classified as *S. marcenscens* by genomic analyses before MLST analysis.

AntiSMASH analysis also revealed that GBS19 produces several antimicrobial compounds that inhibit various pathogenic microorganisms and enhance soil antipathogenic potential (Yegen & Heitefuß, 1970). These protein-encoding genes can be used in recombinant DNA technology for large-scale production and formulation.

*S. sarumanii* GBS19 strain possesses the genetic foundation necessary to activate appropriate metabolic pathway by encoding enzymes. The genes found in the *S. sarumanii* GBS19 strain are linked to degradation processes. We found that the *S. sarumanii* GBS19 strain expresses various genes related to biodegradation and metabolic processes in multiple pathways. Predictive metabolomics analysis revealed that GBS19 exhibits a variety of PVF-mediated signalling properties, which can be linked to cellular physiology and increasing of virulence. The genomic data highlighted the presence of genes encoding toxins, EPS, adhesion factors, and other compounds that are involved in PVF-mediated signalling pathways. These pathways play a significant role in bacterial virulence, colonization, and competition within soil microflora (Kretsch et al., 2021).

Our findings indicated that the *S. sarumanii* GBS19 strain can metabolize some of the active components of tested pesticides to lower quantities and use them as a carbon source. Meanwhile, during the LC-MS analysis, the active substance content of some pesticides increased further after interaction with bacteria. These results should be expanded and confirmed in new studies. However, the results suggest that the combined use of GBS19 at doses below the recommended pesticide concentration can create a synergistic effect that is effective as the recommended dose.

In the pesticide residue degradation analyses, the initial concentration of 11 out of 25 active substances degraded in the solutions in which the bacteria and pesticide were administrated and incubated for 72 h. As is well known, when plants are exposed to pesticides, the chemicals start to degrade from the initial day; however, when they are prepared at the administration dose and preserved in a closed tube until analyses, the degradation rates have shown differentiation on the plant. Pesticide degradation depends on many factors, such as the pH of the application solution and the water solubility coefficient of the active substance, as well as other plant-related factors. On account of the pesticide *K_ow_* values = indicating relationship between lipophilicity and hydrophilicity = directly affect the recovery values in pesticide analyses, the waiting period (72 hours) for some pesticides causes different results depending on the initial concentration. Since the tested bacteria use the main active pesticide molecule as a carbon source, they produce metabolites that change the pH and dissolution coefficient of the solution (water+pesticide). As a result of degradation in the amount of pesticide, it is anticipated that this situation will cause some pesticides to give a higher recovery and response rate, which was detected in the LC-MS analysis (Supplementary table S4). We determined these changes in LC-MS analysis when bacteria were encountered in the pesticide solution. In this study, we noticed that when identifying bacteria at the species level, not only genomic analyses alone but also multilocus sequencing analysis should be performed. When genomic analyses were performed by searching the contig sequences of the bacteria, it was determined that the strain was classified as *Serratia marcescens*, but in subsequent analyses, it was also closely related to *S. nematodiphila*. Based on this finding, it was necessary to expand the analyses and examine allelic variations using MLST analysis. We confirmed that this strain was identified as *S. sarumanii*.

This study demonstrated for the first time that the genetic structure of *S. sarumanii* can produce enzymes that degrade various active compounds. The finding reveals that the potential of the strain owing to its biodegradation property can be integrated into pest management and chemical practice to reduce environmental pollution. Therefore, our study could be a pioneer study of this strain belonging to *S. sarumanii* in purpose of biodegradation. A comprehensive understanding of the enzymes involved in these metabolic pathways linked to biodegradation capacity will enable the application of metabolic engineering or DNA recombinant techniques to use these biomolecules independently of microorganisms. Analyzing the genes involved in pesticide biodegradation is crucial for understanding this process. By understanding the genetic basis of pesticide biodegradation, we can not only trace the evolutionary history of this process but also engineer microorganisms with enhanced degradation capabilities and develop innovative bioremediation strategies for contaminated soils, sediments, and water.

## CONCLUSION

In conclusion, our study demonstrated the potential of the *Serratia sarumanii* GBS19 strain as a promising agent for pesticide biodegradation in agricultural protective applications. By identifying microorganisms capable of breaking down commonly used pesticides, we not only advance environmental conservation efforts and highlight the importance of maintaining the ecosystem balance. The strain’s ability to degrade certain pesticide compounds while enhancing the efficacy of others suggests a dual benefit to sustainable agriculture. Furthermore, the genomic insights gained from this research pave the way for innovative bioremediation applications based on recombinant DNA technology. Overall, these findings underscore the critical role that microbial interventions can play in improving agricultural practices and minimizing environmental impacts.

## CREDIT AUTHORSHIP CONTRIBUTION STATEMENT

GA: Formal analysis; Investigation; Methodology. ÖB: Conceptualization, Data curation; Formal analysis; Funding acquisition; Investigation; Methodology; Project administration; Resources; Software; Supervision; Validation; Visualization; Writing - original draft; Writing - review & editing. RSS: Methodology; Data curation; Funding acquisition; Project administration; Writing - original draft; Writing - review & editing. GÖ and HÖ: Formal analysis; Investigation; Methodology; Data curation. AC: Formal analysis; Methodology.

## DATA AVAILABILITY STATEMENT

The raw sequence data of this study are openly available in NCBI database (NCBI accession: PRJNA784190). The data that supports the findings of this study are available in the supplementary material of this article.

## ACKNOWLEDGEMENTS

The authors wish to thank Nejmettin Kaya, M.Sc. (Marmaris Agriculture District Manager depending on Turkish Ministy of Agriculture and Forestry) for providing pesticides tested in this study.

## ETHICS APPROVAL STATEMENT

Not applicable

## PATIENT CONSENT STATEMENT

Not applicable

## PERMISSION TO REPRODUCE MATERIAL FROM OTHER SOURCES

Not applicable

## CLINICAL TRIAL REGISTRATION

Not applicable

## SUPPLEMENTARY MATERIAL

**Supplementary Table S1.** List of the commercial pesticides tested for pesticide biodegradation according to their active constituents and chemical components.

**Supplementary Table S2.** The media contents used for testing the pesticide biodegradation ability of bacteria.

**Supplementary Table S3.** The pesticide list linked to their conventional agricultural application that were tested using LC-MS analysis in our assays.

**Supplementary Table S4. a)** Pesticide information used in LC-MS analysis, **b)** Information on the LC-MS/MS device used in chromatographic analysis, **c)** Retention time and flow parameters, **d)** Information on the GC-MS device used in chromatographic analyses, **e)** Information on pesticides used in physical and chromatographic analyses, **f)** LC-MS analysis results.

**Supplementary Table S5.** Biochemical tests on isolated bacterial colony.

**Supplementary Table S6.** Degradation pathways associated with genes detected in *S. sarmanuii* GBS19 strain.

**Supplementary Table S7.** Xenobiotic degradation pathways associated with genes detected in *S. sarumanii* GBS19 strain.

**Supplementary Table S8.** Genes encoded at key points of degradation pathways associated with genes of *S. sarumanii* GBS19 strain.

**Supplementary Data S1.** Organism overview for proteobacteria *Serratia* strain GBS19.

**Supplementary Data S2.** Compound calibration report related LC-MS analysis.

**Supplementary Data S3.** Genome annotation data.

**Supplementary Data S4.** Genome annotation data (gbk. file).

**Supplementary Data S5**. Type (strain) genome server results.

**Supplementary Data S6**. Ribosomal MLST results.

**Supplementary Data S7**. Genomic MLST results.

**Supplementary Data S8.** KEGG pathway maps of *S. sarumanii*.

**Supplementary Data S9**. AntiSMASH results.

**Supplementary Figure S1.** Visual appearance of **a)** pure bacterial colonies and **b)** microscopic image of Gram staining.

**Supplementary Figure S2.** Pie chart of metabolic pathways and their related feature counts.

**Supplementary Figure S3.** Pathogen IslandViewer 4.0 output of GBS19.

